# Androgen Receptor-Induced Integrin α6β1 and Adhesion to Laminin Promotes Survival and Drug Resistance in Castration-Resistant Prostate Cancer through BNIP3

**DOI:** 10.1101/349050

**Authors:** Eric A. Nollet, Sourik S. Ganguly, Veronique V. Schulz, Anne Cress, Cindy K. Miranti

## Abstract

Although castration-resistant prostate cancers no longer respond to anti-androgen therapies, the androgen receptor (AR) is still required to promote tumor survival. However, the signaling pathways downstream of AR that promote this survival are not well known. We recently identified an AR-dependent survival pathway whereby AR induction of integrin α6β1 and adhesion to laminin activates NF-κB/RelA signaling and Bcl-xL. This pathway acts in parallel with the PI3K/Akt pathway in Pten-null tumor cells such that combined inhibition of both PI3K and integrin α6β1 is required to kill tumor cells adherent to laminin. However, PTEN-null castration-resistant tumors were not effectively inhibited by this combination. We discovered that BNIP3, a hypoxia-induced BH3-only, pro-mitophagic Bcl2 family member, is induced by androgen in castration-resistant cells through integrin α6β1 signaling to HIF1α. Furthermore, castration-resistant cells adherent to laminin were much more efficient at inducing autophagy in response to androgen. Androgen blocked the ability of the PI3K inhibitor PX-866 to kill castration-resistant tumors, but this was reversed by loss of BNIP3. Although BNIP3 was dispensable for androgen-induced autophagy, its mitophagy function was required for BNIP3 to promote resistance to PI3K inhibition. Thus, adhesion to laminin triggers signaling through AR/α6β1/HIF1α in castration-resistant cells to drive the expression of BNIP3 and cooperates with AR/α6β1-mediated autophagy, both of which contribute to PI3K resistance through induction of mitophagy.

## INTRODUCTION

Androgen deprivation therapy is the standard of care for men with advanced or metastatic prostate cancer. However, like most cancer therapies, patients will eventually relapse and no longer respond to the therapy. This is even true for 2^nd^ generation therapies that target the androgen pathway (1). Thus, there is a need to understand the mechanisms that underlie anti-androgen therapy resistance, also referred to as castration-resistance. Many mechanisms have been proposed, but the common underlying theme is that the target of anti-androgen therapy, the androgen receptor (AR), remains functionally intact through amplification, splice variants, and supplementary signaling pathways (2). What is not well defined are the signaling pathways that specifically lie downstream of AR that allow survival of the tumor cells during and after therapy resistance. Identification of these pathways will be necessary to design new therapeutic approaches to combat castration-resistant disease.

Loss of expression of the tumor suppressor, Pten, is a common occurrence in prostate cancer, especially in castration-resistant and metastatic disease. Loss of PTEN occurs in 40 - 60% of castration-resistant prostate cancers (3), and is one of the few markers that predicts for poor outcome. One consequence is constitutive activation of the PI3K/Akt survival pathway (4). Thus, it was expected that pharmacological inhibitors of this pathway and downstream effectors, such as mTor, would be effective therapies. While these drugs can repress cancer cell growth *in vitro* and in xenograft transplants, as single agents they have not been effective in patients (5). Thus, there are likely additional survival pathways operating *in vivo*.

We previously identified an AR-dependent survival pathway in metastatic prostate cancer cell lines (6). In prostate cancer, integrin β4 is downregulated while integrin α6 is upregulated causing a shift in α6 pairing with β1 (7). We demonstrated that AR directly induces the expression of integrin α6β1 such that adhesion to its ligand, laminin, activates NF-κB/RelA signaling and induction of the anti-apoptotic protein, Bcl-xL. This pathway acts in parallel with the PI3K/Akt pathway in Pten null tumor cells such that combined inhibition of both PI3K and integrin α6β1 is required to effectively kill these cells *in vitro* when they are adherent to laminin (6). Because laminin is a major extracellular matrix component of the lymph node and bone microenvironment (8,9), this pathway is likely to be a very important mechanism for escaping drug therapy (10).

Androgen deprivation therapy or chemical castration is known to induce a hypoxic environment within the prostate (11,12). The hypoxic response, mediated by HIF1α/HIF2α, is a well-established mechanism that protects tumor cells from low oxygen environments. Elevated levels of HIF1α and HIF2α have been reported in castration-resistant and metastatic prostate cancer (13). Furthermore, hypoxia directly stimulates the expression of integrin α6 and β1 (12,14). Thus, hypoxia-induced integrin α6β1 expression may provide an additional mechanism for prostate cancer survival under androgen-deprivation conditions.

HIF1α/HIF2α mediate their survival effects under hypoxic conditions in part through direct transcriptional induction of target genes. One of those targets is BNIP3 (15,16). BNIP3 was reportedly elevated in castration-resistant cells after passage *in vivo* in castrated mice (17), and elevated BNIP3 expression in human prostate cancers predicts for poor outcome (18,19). BNIP3, a BH3-only member of the Bcl2 family, is localized on mitochondria and reportedly induces apoptosis, autophagy, or mitophagy (20). The mechanism by which it induces apoptosis is not completely clear, but seems to be primarily a result of excessive overexpression (21). It is proposed to control autophagy through its ability to interact with Bcl-xL, thus removing the negative constraint on the pro-autophagy protein, Beclin (22). BNIP3 contains an LC3-interacting region (LIR), which binds LC3 on nascent autophagy membranes and thereby recruits mitochondria into autophagosomes (23). BNIP3 association with Bcl-xL also enhances the affinity of BNIP3 for LC3 (23). Genetic evidence demonstrates a role for BNIP3, and its homologue NIX/BNIP3L, in regulating mitochondrial clearance. BNIP3-/- mice accumulate defective mitochondria in hepatocytes and acquire phenotypes of metabolic dysfunction (24), while NIX/BNIP3L-/- mice cannot clear mitochondria out of maturing erythrocytes (25).

In this study, we tested the hypothesis that BNIP3 is the control point that links the AR/integrin α6β1 pathway and the hypoxia pathway to promote the survival of castration-resistant prostate cancer.

## Materials and Methods

### Cell Culture

Tissue culture plates were coated with 10 μg/mL mouse laminin (Gibco: 23017-015) in Ca+/Mg+-free (CMF) PBS overnight at 4° C. LNCaP cells, purchased from ATCC, and C4-2 cells, obtained directly from Dr. Robert Sikes (17), were both validated by autosomal STR analysis in our Genomics core. Cells were discarded after 30 passages and replaced with low-passage cells to maintain consistency throughout the study. Cells were routinely passaged and maintained on laminin in RPMI supplemented with 10% FBS, 1 mM sodium pyruvate, 2 mM glutamine, 0.3% glucose, 10 mM HEPES, and 30 U/mL Pen/Strep. HEK293FT cells (Clontech), used for lentivirus generation, and Phoenix-AMPO cells (National Gene Vector Biorepository), used for retrovirus generation, were maintained in DMEM supplemented with 10% HIFBS and 30 U/mL Pen/Strep.

### Androgen and drug treatments

Laminin-coated plates were blocked with 1% BSA in CMF PBS at 37°C for 1 hour prior to plating cells. Due to differential proliferation rates, C4-2 were plated at 30,000 per cm^2^ while LNCaP were plated at 60,000 per cm^2^, to ensure similar cell densities by the end of the experiment. After 24 hours, the media was changed to starvation media (Phenol red-free RPMI supplemented with 0.1% charcoal-stripped serum, 1 mM sodium pyruvate, 2 mM glutamine, 0.3% glucose, 10 mM HEPES, and 30 U/mL Pen/Strep). When needed for shRNA induction, doxycycline was added to a final concentration of 100 ng/mL. After 48 hours, cells were stimulated with 10nM R1881 (synthetic androgen) or ethanol vehicle and re-spiked with doxycycline when used. Twenty-four hours later cells were treated with 20-500nM PX-866 (26) or DMSO vehicle for 48 hours.

### Cell death assay

Cell death was measured after 48 hours after PX-866 treatment. Attached and floating cells were collected, pooled, and stained with Trypan blue. For each individual experiment 3 wells per condition and at least 3 grids per well were counted on a hemocytometer.

### Integrin blocking

Cells were trypsinized, spun down, and counted. Suspended cells at 30,000 cells/mL in starvation media were treated with 10 μg GoH3 anti-integrin α6 antibody or vehicle. The cells were then plated on laminin-coated 12-well plates at a final concentration of 30,000 cells/cm^2^. After 48 hours, cells were stimulated with 10nM R1881 or DMSO vehicle. Twenty-four hours later cells were treated with 500nM PX-866 (26) or DMSO vehicle. Cell death was measured after 48 hours after PX-866 treatment as described above.

### qRT PCR

RNA was isolated using Trizol and lithium chloride precipitation or using the RNEasy kit from Qiagen. RNA was reverse transcribed using MuLV reverse transcriptase (New England Biolabs) with a mix of random hexamers and polyT primers. cDNA was amplified using FastStar Universal SYBR Green Master (Rox, Roche) in the Applied Biosystems 7500 RT PCR System. Levels of target mRNAs were normalized to 18S ribosomal RNA. List of primers is in Supplementary Table S1. For protein synthesis inhibition, cycloheximide in ethanol was added to a final concentration of 10 μg/mL coincidental with R1881 treatment.

### Immunoblotting

30-60 μg of protein were loaded in precast tris-glycine gels from Invitrogen and transferred to a PVDF membrane. Membranes were blocked with 5% BSA TBST and all antibodies were diluted in 5% BSA TBST. Primary antibodies were detected using HRP conjugated secondary antibodies in chemiluminescent solution using the Quantity One imaging software on a Bio-Rad Gel Docking system. <u>*Primary antibodies:*</u> Rabbit mAb BNIP3 (EPR4034) (Abcam), rabbit anti-P-AktSer473 (Cell Signaling Technology), rabbit mAb P-Akt308 (C31E5E) (Cell Signaling Technology), rabbit mAb Akt (pan) (C67E7) (Cell Signaling Technology), mouse mAb GAPDH (Millipore), mouse mAb Histone H3 (96C10) (Cell Signaling Technology), rat anti-integrin α6 (GoH3) (BD Pharmingen), mouse mAb AR (441)(Santa Cruz), mouse anti-alpha tubulin (Sigma-Aldrich), mouse anti-HIF1α (BD Pharmingen, and rabbit anti-integrin α6 (AA6A, A6NT) obtained from Dr. Anne Cress (27,28), University of Arizona.

### Virus generation and infection

For lentiviral constructs, 5 × 10^6^ HEK293FT cells, and for retroviral constructs, 5 × 10^6^ Phoenix-AMPO cells, were plated in DMEM with 10% HIFBS in a T75 flask that was coated with 2 μg/mL Poly-D-lysine in PBS and left overnight at 37 °C. The following day the cells were transfected in Opti-MEM using Lipofectamine2000 with 5 μg pLP1, 5 μg pLP2, 5 μg pVSV-G, and 5 μg of target construct. After 24 hours the media was changed to RPMI with 10% HIFBS and no antibiotics and returned to 37 °C for 48 hours. Floating cells were spun out of solution, and the supernatant was filtered through a 0.45 μm filter to remove debris. Polybrene was added to a final concentration of 5 μg/mL to the filtered media and this was added to target cells and incubated at 32°C. Six hours after virus treatment, cells were washed and returned to normal media and subjected to antibiotic selection.

### siRNA

C4-2 cells were plated on laminin at a density of 30,000 cells/cm^2^. Twenty-four hours later cells were transfected in antibiotic-free starvation media (Phenol red-free RPMI supplemented with 0.1% charcoal-stripped serum, 1 mM sodium pyruvate, 2 mM glutamine, 0.3% glucose, 10 mM HEPES) using siLentFect (BioRad) and siRNA at a final concentration of 20 nM. Twenty-four hours later, the medium was changed to fresh starvation medium. siRNAs used include: Two different HIF1α siRNAs (J-004018-07 and J-004018-08, Dharmacon), one integrin α6 siRNA (5’- CGAGAAGGAAATCAAGACAAA-3’), and a non-targeting siRNA (D-001206-14, Dharmacon).

Doxycycline-inducible and constitutive shRNAs: Doxycycline-inducible shRNA plasmids targeting integrin α6 and BNIP3 were generated by sub-cloning shRNA sequences into a Tet-inducible lentiviral vector, EZ-Tet-pLKO-Puro (49), available through Addgene (#85966). After infection of C4-2 cells, cells were selected in 2 μg/mL puromycin and single cells were isolated to generate clonal lines. During experiments the cells were treated with doxycycline at a final concentration of 100 ng/mL to induce shRNA expression. Targeting sequences are in Supplementary Table S2. The same integrin α6 targeting shRNA sequence was also cloned into pLKO.1 Puro (gift from Bob Weinberg (Addgene plasmid # 8453)) (29) and used to generate a stable cell line. A scrambled non-targeting shRNA was used as vector control.

### GFP-LC3 quantification

LNCaP and C4-2 cells were infected with retrovirus containing pBABEPuro GFP-LC3 (gift from Jayanta Debnath (Addgene plasmid # 22405)) (30) and selected with 2 μg/mL puromycin. Cells were plated on laminin-coated glass coverslips, and treated for 24 hours with 10nM R1881 or vehicle (ethanol), and then for the last 2 hours with or without Bafilomycin A1 and mounted on glass slides. Using a 60X objective, 25 fields were selected per condition per experiment and the number of puncta counted. Puncta were considered positive if they were 10 standard deviations brighter than background fluorescence and within the size range of an HBSS-treated positive control.

### LC3-II immunoblot quantification

LNCaP and C4-2 were plated on laminin in 6 cm plates. After 24 hours, the media was changed to starvation media, and 48 hours later R1881 was added to a final concentration of 10 nM. Bafilomycin A1 (100 ng/mL) or vehicle control DMSO was added 2 hours prior to lysis 24 hours after R1881 treatment. Blot density was measured using ImageJ software. The density of LC3-II was normalized to tubulin density in the same lane. The ratio of the first lane was set to one, and subsequent lanes are relative to the first (control) lane. Ratios were calculated in 3 separate experiments.

### BNIP3 re-expression

A plasmid, pENTR223-BNIP3 (HsCD00366502) containing the BNIP3 cDNA, was obtained from the Harvard PlasmID repository (31). Site-directed mutagenesis was used to first generate a stop codon in pENTR223-BNIP3 vector. pENTR223-BNIP3 underwent two more rounds of site directed mutagenesis to eliminate the LC3-interacting region (LIR) to generate pENTR223-BNIP3 ΔLIR in which amino acids W18 and L21 were converted to alanine. Mutagenesis primers are listed in Supplementary Table S3. These mutations are sufficient to eliminate BNIP3 interaction with LC3 (23,32). BNIP3 WT and ΔLIR were each recombined into a Tet-inducible lentiviral vector, pLenti CMVTight Neo DEST (gift from Eric Campeau (Addgene plasmid # 26432) (33), using LR recombinase to generate the pLenti CMVTight Neo BNIP3 WT and ΔLIR. C4-2 cells selected in 2 μg/mL puromycin and constitutively expressing BNIP3 shRNA targeting the 3’-UTR (SHCLNG-NM_004052, 5’-CCACGTCACTTGTGTTTATT-3’; Sigma-Aldrich) were infected with lentivirus expressing doxycycline-regulated rtTA in pLentiCMV rtTA3 Blast (Addgene #26429; Eric Campau) (33) and further selected in 5 μg/mL blasticidin. An isolated pool expressing rtTA was then infected with pLenti CMVTight Neo BNIP3 WT or ΔLIR lentivirus and further selected in 100 ng/mL G418 (neomycin) to generate a triple antibiotic-resistant pool.

### Mouse studies

Mouse studies were conducted according to an IACUC approved protocol (13-11-033) at Van Andel Institute. A total of 1 × 10^6^ cells in 10 μL DMEM of a C4-2 clonal cell line harboring the doxycycline-inducible shRNA targeting BNIP3 were injected orthotopically into the prostates of 20 6 week-old male nude mice concurrent with castration surgery. Mice were randomly divided into 2 cohorts of 10 each. Immediately after surgery, mice from one group were given water containing 5% sucrose and mice from the other group were given water with 1mg/ml doxycycline (Dox) in 5% sucrose, which was replaced weekly. Mice were sacrificed eight weeks later. Prostate tumors were excised and weighed and assessed by IHC. Ki67 IHC staining was quantified by counting number of positive nuclei in five random fields per sample.

### Immunohistochemistry

Prostate tumors were fixed in 10% neutral buffered formalin, embedded in paraffin blocks, and sectioned. Proteins were probed by IHC using rabbit mAb to cleaved Caspase-3 (Asp175) (5A1E) (Cell Signaling Technology), rabbit mAb BNIP3 (EPR4034) (Abcam), and anti-Ki67 (SP6) (Thermo-Scientific) at 1:00 and incubating overnight at 4°C. Immunostaining was visualized using the DAKO Envision+ Dual link System (DAB+) kit.

## RESULTS

### Integrin α6 and integrin β1 expression positively correlate with castration resistance

The LuCaP tumor series was generated by engrafting and serially passaging human patient-derived (PDX) prostate cancers subcutaneously in mice (34). Castration-resistant variants were generated for some of these lines by passage through castrated mice. Gene expression analysis comparing the parental androgen-sensitive lines to the castration-resistant lines indicated elevated integrin α6 and integrin β1 expression correlated with castration resistance (Fig. 1A).

**Figure 1:**
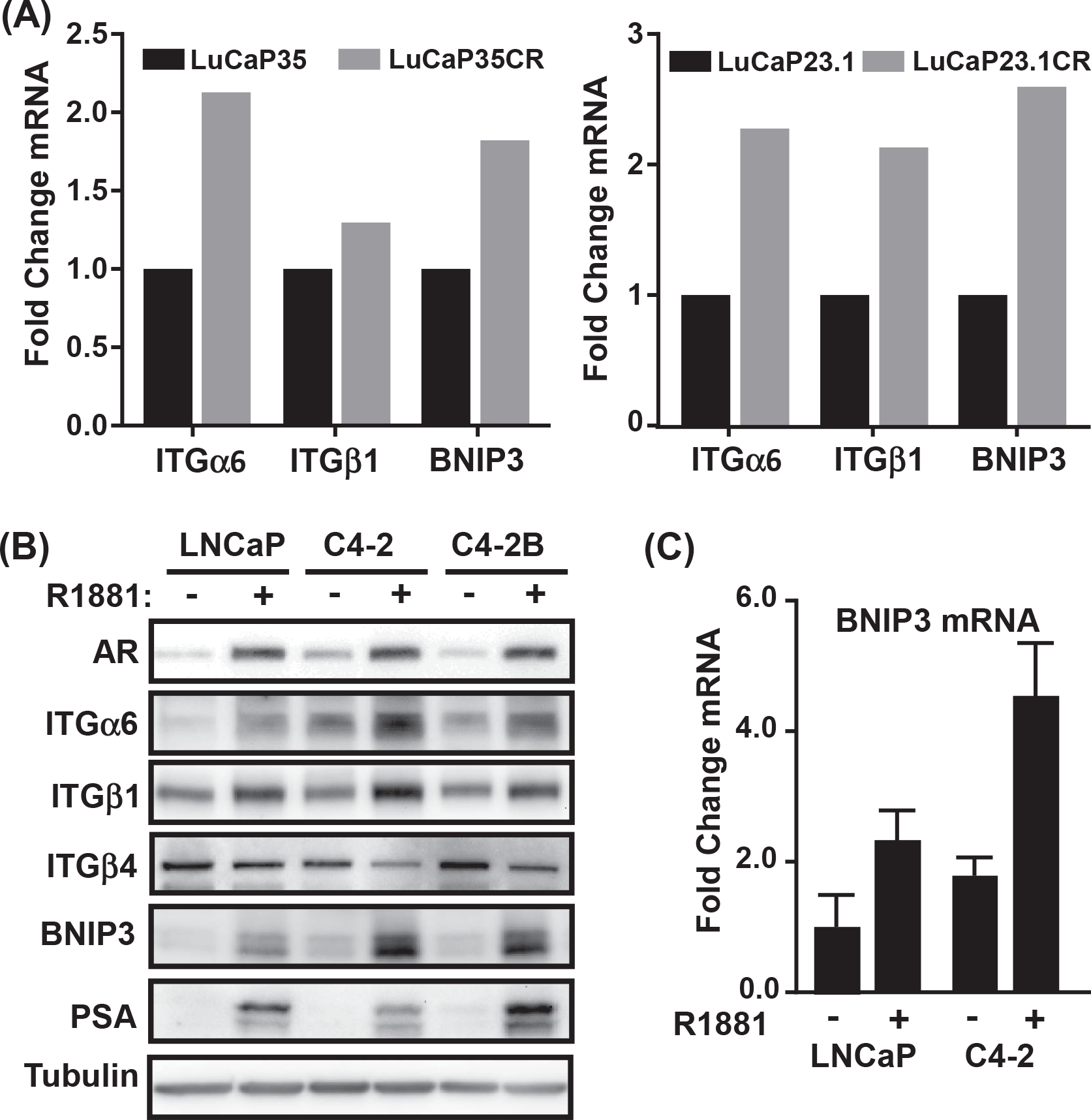
Integrin α6β1 and BNIP3 are elevated in castration-resistant prostate cancer. **(A)** Gene expression of integrin α6 (*ITGα6*), integrin β1 (*ITGβ1*), and *BNIP3* in castration-resistant (CR) human-derived xenograft variants of LuCaP23.1 and LuCaP35 normalized to expression in androgen-sensitive parental tumors. **(B)** Levels of androgen receptor (AR), integrin α6 (ITGα6), integrin β1 (ITGβ1), integrin β4 (ITGβ4), BNIP3, PSA, and tubulin in laminin-adherent LNCaP, C4-2, and C4-2B cells with (+) and without (-) stimulation with 10 nM R1881 for 24 h were assessed by immunoblotting. **(C)** Levels of BNIP mRNA in laminin-adherent LNCaP and C4-2 cells with (+) or without (-) stimulation with R1881 for 24 h were measured by qRT-PCR.

The LNCaP prostate cancer cell line was originally isolated from the lymph node of a patient who had not undergone androgen deprivation therapy (35). LNCaP cells express a mutant AR, but are still dependent on androgen for proliferation and survival (36). LNCaP also harbor a frameshift mutation in PTEN, which allows for unrestricted activation of PI3K (37). Castration resistant LNCaP derivatives were generated by serially passaging LNCaP through castrated mice to generate a castration resistant prostate cancer line, C4-2 (38). A C4-2B variant was then derived by passage of C4-2 cells in bone of castration-resistant mice. Basal integrin α6 expression was higher in the castration-resistant C4-2 and C4-2B cells relative to LNCaP, and stimulation of AR by synthetic androgen, R1881, increased both integrin α6 and β1 expression in all cell lines, but to a higher degree in the C4-2 and C4-2B cells (Fig. 1B). Conversely, integrin β4 was decreased by androgen as previously reported (6). Thus, elevated integrin α6β1 expression correlates with progression to castration resistance and is stimulated by androgen.

### BNIP3 expression positively correlates with castration resistance

Analysis of pro-survival proteins in the LuCaP expression profiles identified elevated BNIP3 expression in the castration-resistant variants (Fig. 1A). Microarray data comparing LNCaP and C4-2 cells previously identified significant elevation of BNIP3 mRNA in C4-2 cells (17). Furthermore, elevated BNIP3 expression in human prostate cancers predicts for poor outcome (18,19). We similarly observed increased BNIP3 expression in C4-2 and C4-2B cells relative to LNCaP, at both the transcript and protein level (Fig. 1B, C). Furthermore, we found that BNIP3 expression increases after androgen stimulation (Fig. 1B, C). Thus, BNIP3, like integrin α6β1, correlates with progression to castration resistance and is induced by androgen.

### Androgen stimulates BNIP3 expression through Integrin α6 and HIF1α

BNIP3 is a classic hypoxia target, transcriptionally induced directly by HIF1α (15,16). Both HIF1α and HIF2α are elevated in the two castration-resistant variant lines (Supplementary Fig. S1A). To determine if AR can directly induce BNIP3 transcription, we compared the kinetics of BNIP3 mRNA induction to the known AR target gene, PSA. While PSA mRNA increased immediately (within 3-6 hrs), BNIP3 transcript did not significantly increase until 18 hours after R1881 stimulation (Fig. 2A). Furthermore, blocking protein synthesis with cyclohexamide prevented R1881 from increasing BNIP3 transcription (Fig. 2B) indicating BNIP3 is not a direct target of AR. To determine whether androgen stimulation of HIF1α or integrin α6 was required for BNIP3 expression, we monitored the localization and induction of HIF1α, integrin α6, and AR over time following androgen stimulation. In the first 6 hours, AR shifted into the nucleus and the direct targets of AR transcription, PSA and integrin α6, increased as expected (Fig. 2C). C4-2 cells constitutively express HIF1α, and while androgen did not significantly increase its expression, it did induce nuclear accumulation at 12 hours. BNIP3 protein did not increase until after 12 hours (Fig. 2C), suggesting that androgen induction of integrin α6 and HIF1α nuclear localization is required prior to BNIP3 induction.

**Figure 2:**
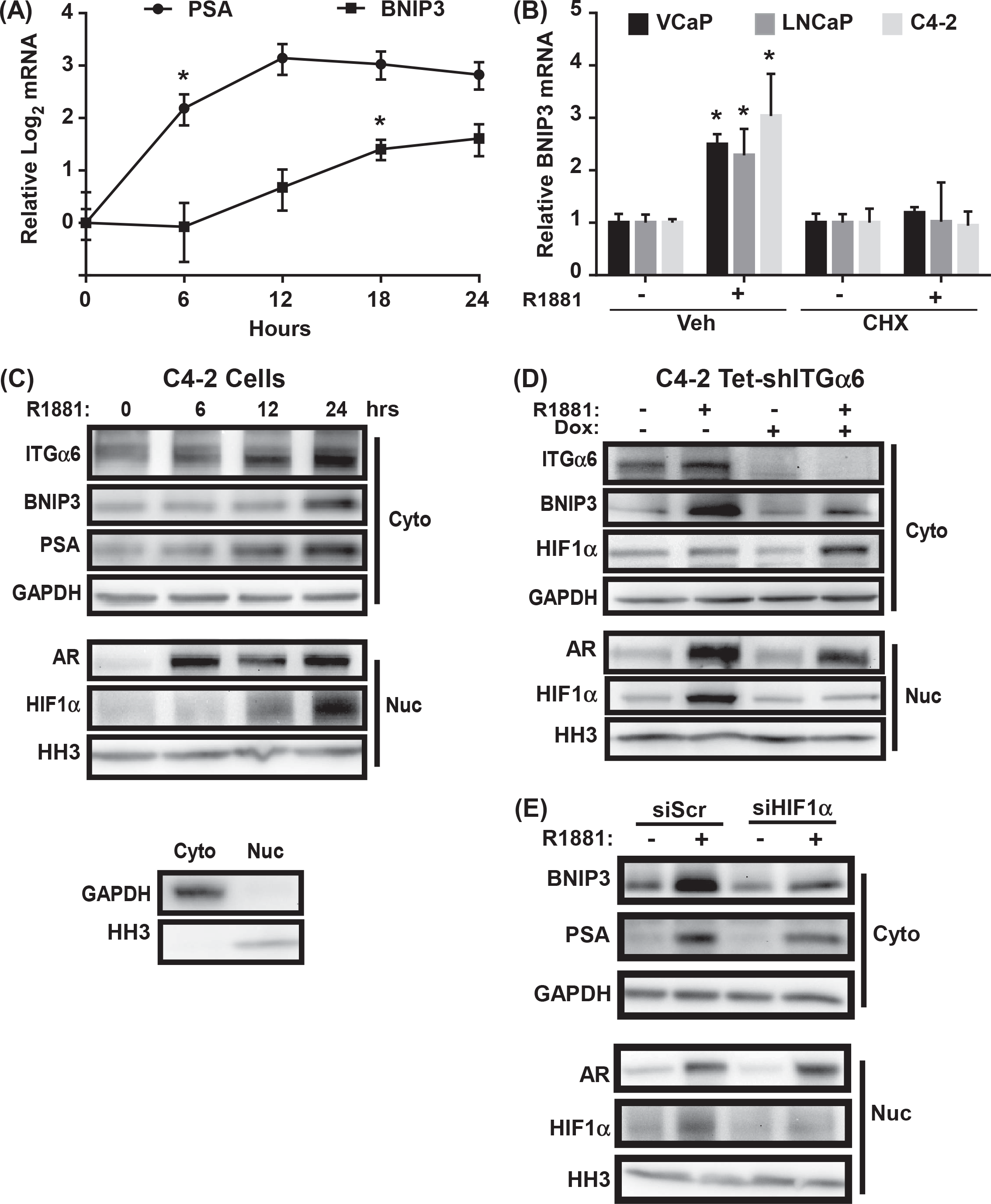
Androgen indirectly induces BNIP3 through integrin α6β1 and HIF1α. **(A)** Levels of PSA and BNIP3 mRNA in laminin-adherent C4-2 cells over a 24 h time course following stimulation with 10 nM R1881 were measured by qRT-PCR and normalized to time 0. * *p* < 0.05, n=3; error bars = SD. **(B)** Levels of BNIP3 mRNA in laminin-adherent VCaP, LNCaP, and C4-2 cells stimulated with (+) or without (-) 10 nM R1881 in the absence (Veh) or presence of 10 μg/mL cycloheximide (CHX) for 24 hours prior to lysis. Expression is relative to control. * *p* < 0.05, n=3, error bars = SD. **(C)** Levels of integrin α6 (ITGα6), BNIP3, PSA, and GAPDH in the cytosol (Cyto) and androgen receptor (AR), HIF1α, and histone 3 (HH3) in the nucleus (Nuc) of laminin-adherent C4-2 cells stimulated with 10 nM R1881 for time course of 24 h were measured by immunoblotting. **(D)** Tet-inducible shRNA targeting integrin α6 (Tet-shITGα6) was stably expressed in C4-2 cells. Following adhesion to laminin, the levels of integrin α6 (ITGα6), BNIP3, PSA, and GAPDH in the cytosol (Cyto) and androgen receptor (AR), HIF1α, and histone 3 (HH3) in the nucleus (Nuc) in response to 10 nM R1881 with (+) or without (-) 100 ng/mL doxycyclicne (DOX) to induce shITGα6 were assessed by immunoblotting. **(E)** C4-2 cells were transiently transfected with HIF1α-specific siRNA. Forty-eight hours later, following adhesion to laminin, the levels of BNIP3, PSA, and GAPDH in the cytosol (Cyto) and AR, HIF1α, and histone H3 (HH3) in the nucleus (Nuc) following R1881 stimulation for 24 h were assessed by immunoblotting.

Inhibition of integrin α6 expression by a Tet-inducible shRNA, not only prevented BNIP3 induction in response to androgen, but also blocked HIF1α nuclear localization (Fig. 2D). Inhibition of HIF1α by two different siRNAs prevented androgen-induced BNIP3 mRNA and protein expression (Fig. 2E, Supplemental Fig. S1B,C), but had no impact on integrin α6 expression (Supplemental Fig. S1C). Thus, integrin α6 is required for androgen to stimulate HIF1α nuclear translocation to induce BNIP3 expression.

### C4-2 cells have decreased sensitivity to PX-866

We previously demonstrated that inhibition of both PI3K and integrin α6 is required to effectively kill AR-expressing cells adherent to laminin (6). To determine if this is also true for castration resistant cells, LNCaP and C4-2 cells were first treated with PX-866, a class I-selective PI3K inhibitor (26). The LD50 for LNCaP was 15 nM, while the C4-2 LD50 was 320 nM, indicating a striking resistance of C4-2 cells to PI3K inhibition (Fig. 3A). Moreover, even at the highest drug concentrations (500 nM), stimulating C4-2 cells with androgen made them completely resistant to PX-866 (Fig. 3A). This level of protection was not observed in LNCaP cells, where androgen did not significantly change the LD50 of PX-866. One possibility is that androgen increases PI3K signaling or reduces the efficacy of PX-866 to block PI3K signaling. However, androgen did not increase Akt activation, nor did it prevent PX-866 from inhibiting Akt (Fig. 3B). Thus, there is something else uniquely mediating androgen resistance in the castration resistant C4-2 cells.

**Figure 3:**
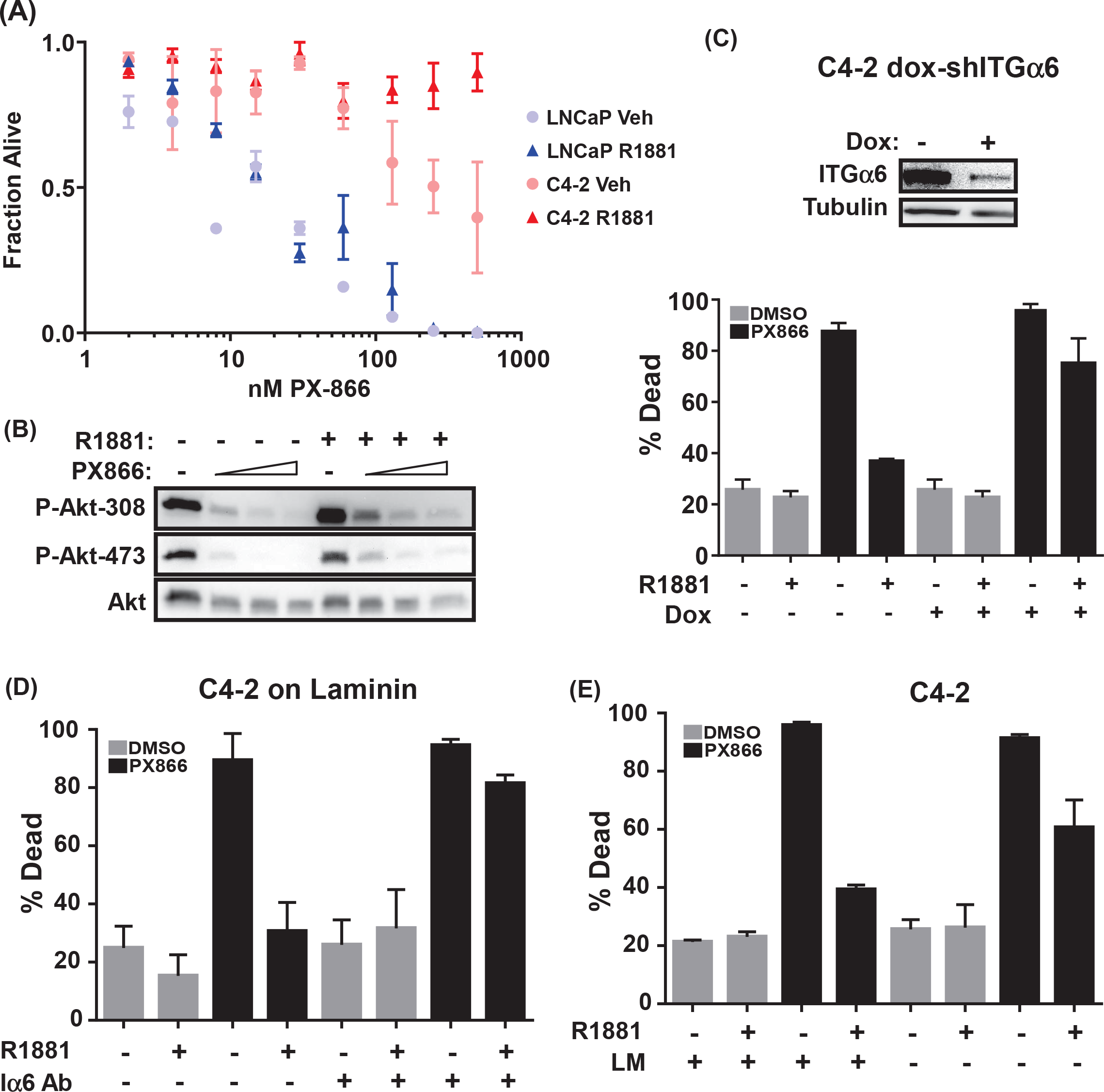
AR confers resistance to PI3K inhibition via integrin α6β1. **(A)** LNCaP and C4-2 cells adherent to laminin were stimulated with (R1881) or without (Veh) 10 nM R1881 for 24 h prior to and in the presence of increasing concentrations of PX-866. Cell viability was measured by trypan blue exclusion 48 h later. n=3, error bars = SD. **(B)** Laminin-adherent C4-2 cells were stimulated with (+) or without (-) 10 nM R1881 for 24 h prior to and in the presence of 0, 100, 200, and 400 nM PX-866. Levels of activated Akt (P-Akt-308; P-Akt-473) and total Akt were measured by immunoblotting 48 h later. **(C)** Levels of integrin α6 (ITGα6) and tubulin in laminin-adherent C4-2 cells stably expressing Tet-inducible shRNA targeting integrin α6 (Tet-shITGα6) were measured by immunoblotting following induction by 100 ng/mL doxycycline (DOX). Laminin-adherent C4-2 cells were stimulated with (+) or without (-) 10 nM R1881 and with (+) or without (-) 100 nM doxycycline (DOX) for 24 h prior to and in the absence (DMSO) or presence (PX866) of 500 nM PX-866. Cell viability was assessed by trypan blue exclusion 48 h later. n=3, error bars = SD. **(D)** C4-2 cells were treated with 10 μg/mL integrin α6 GoH3 blocking antibody (Iα6 Ab, +) or without (Iα6 Ab, -) prior to plating on laminin. Cells were then stimulated with (+) or without (-) 10 nM R1881 24 h prior to and in the presence (PX866) or absence (DMSO) of 500 nM PX-866. Cell viability was measured by trypan blue exclusion 48 h later. n=3, error bars = SD. **(E)** C4-2 cells plated on laminin (LM, +) or on plastic (LM, -) were stimulated with (+) or without (-) 10 nM R1881 for 24 h prior to and in the presence (PX866) or absence (DMSO) of 500 nM PX-866. Cell viability was measured by trypan blue exclusion 48 h later. n=3, error bars = SD.

### Integrin α6 and Bnip3 protect C4-2 cells from PI3K inhibition

To determine whether integrin α6 is required to protect C4-2 from PX-866, we blocked integrin α6 expression with shRNA, inhibited integrin α6 adhesion to laminin with blocking antibody, or plated cells on plastic without laminin. Blocking integrin α6 function completely blocked the ability of androgen to protect cells from high levels (500nM) of PX-866 (Fig. 3C-E). Thus, integrin α6-mediated adhesion to laminin is required for androgen to protect C4-2 cells from PI3K inhibition.

To determine whether BNIP3 is a downstream target of integrin α6-mediated survival, BNIP3 expression was inhibited with two different Tet-inducible shRNAs. Without BNIP3, androgen stimulation could not protect the cells from death induced by PX-866 (Fig. 4A, B). PX-866 itself did not inhibit BNIP3 expression and the shRNA effectively knocked down BNIP3 (Fig. 4C). Thus, BNIP3 is required to protect C4-2 cells from PI3K inhibition.

**Figure 4:**
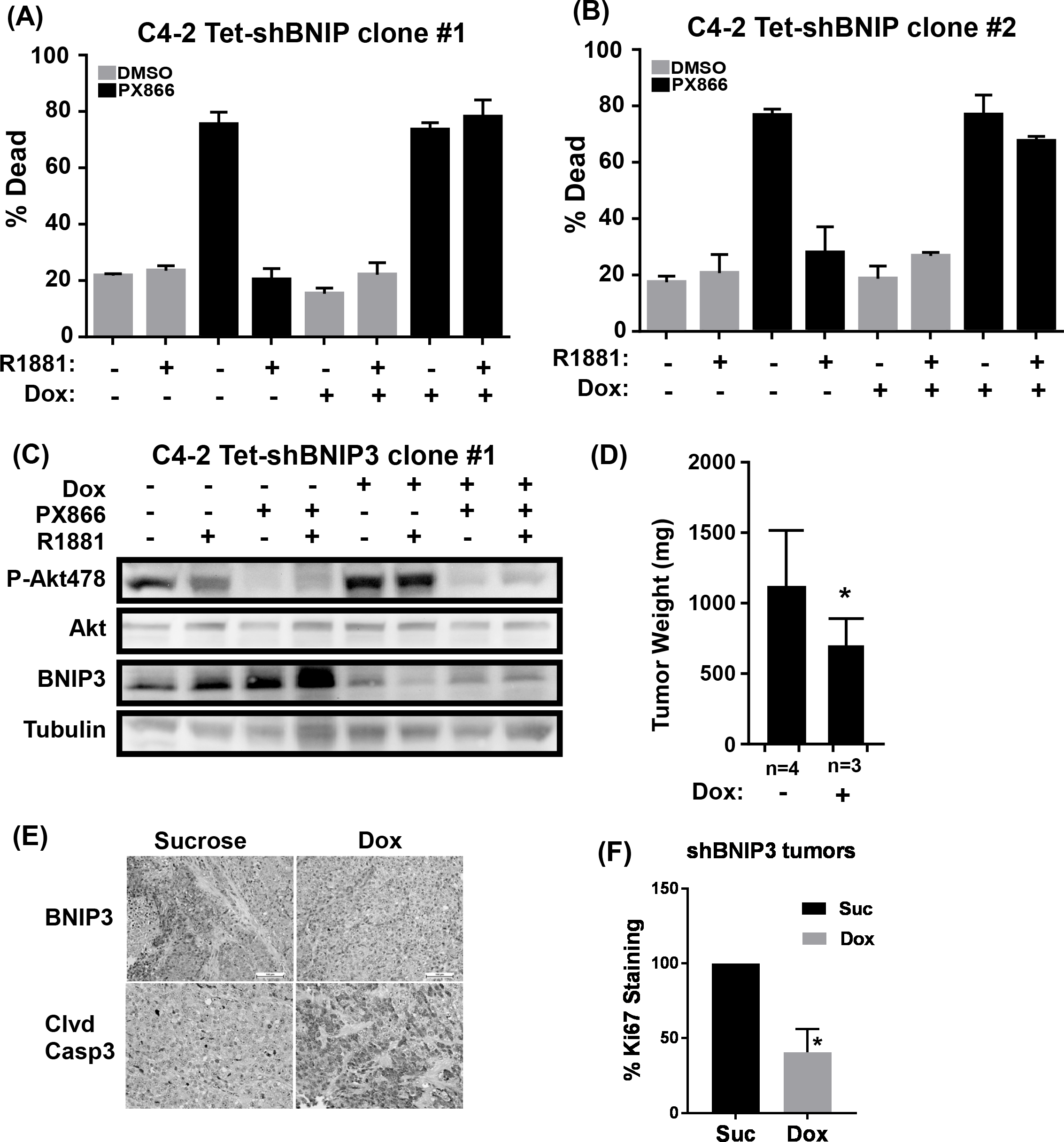
AR confers resistance to PI3K inhibition via BNIP3. **(A, B)** Two different Tet-inducible BNIP3 shRNAs (Tet-shBNIP3) were stably introduced into C4-2 cells. Cell viability of laminin-adherent stimulated concurrently with (+) or without (−) 10 nM R1881 and with (+) or without (−) 100 ng/mL doxycycline (DOX) 24 h prior to and in the absence (DMSO) or presence of 500 nM PX-866 was assess by trypan blue exclusion 48 h later. n=3, error bars = SD. **(C)** Levels of activated Akt (P-Akt478), total Akt, BNIP3, and tubulin in (A) were assessed by immunoblotting. **(D)** C4-2 cells stably expressing Tet-inducible BNIP3 shRNA (Tet-shBNIP3) were injected orthotopically into prostates of castrated mice. Mice were fed sucrose (DOX, -) or sucrose with doxycycline (DOX, +) for 8 weeks and then tumors were harvested and weighed. * *p* < 0.10, n=3, error bars = SD. **(E)** Representative tumor tissues from mice in (D) were assessed for expression of BNIP3 and cleaved caspase 3 (Clvd Casp3). **(F)** Representative tumor tissues from mice in were assessed for expression of Ki67 by IHC and quantified. * *p* = 0.0027, n=3, error bar = SD.

### BNIP3 promotes castration-resistant tumor growth and survival

A total of 1 × 10^6^ C4-2 cells harboring the Tet-inducible BNIP3 shRNA were injected orthotopically into the prostate glands of 20 castrated male nude mice. Half of the mice, 10 per cohort, were fed doxycycline in 5% sucrose in their drinking water, the other half only received sucrose. Eight weeks later, mice were sacrificed and glands, tumors, and the proximal lymph nodes were collected and fixed, and the gross tumor weight recorded. The tumors from mice treated with doxycycline (i.e. loss of BNIP3) were about twice as small as the controls (Fig. 4D). We verified BNIP3 expression was decreased in the doxycycline-treated tumors (Fig. 4E). Loss of BNIP3 reduced proliferation, as measured by Ki67 (Fig. 4F) and increased cell death, as measured by cleaved caspase 3 IHC (Fig. 4E), both likely contributing to the decrease in tumor volume. Thus, BNIP3 expression is contributing to castration-resistant tumor growth and survival.

### Androgen increases autophagy in CRPC cells on laminin via integrin α6, independent of BNIP3

BNIP3 could be promoting CRPC survival through autophagy and/or mitophagy (20), and either mechanism requires an overall increase in macroautophagy in castration-resistant cells. Furthermore, because BNIP3 induction requires androgen, then androgen stimulation should induce autophagy if BNIP3 is involved. We first measured autophagy by examining the accumulation of GFP-labelled LC3B puncta on autophagosomes in LNCaP and C4-2 cells adherent to laminin following androgen stimulation with and without bafilomycin A1 (39). At steady state, both LNCaP and C4-2 have similar numbers of autophagosomes, and this is not significantly changed by the addition of only androgen. However, treatment with bafilomycin, to stop lysosomal turnover of autophagosomes and LC3-II degradation, reveals that autophagosome accumulation occurs at a significantly higher extent in C4-2 cells treated with androgen (Fig. 5A,B). Thus, the CRPC cells more robustly activate autophagic turnover in response to androgen.

**Figure 5:**
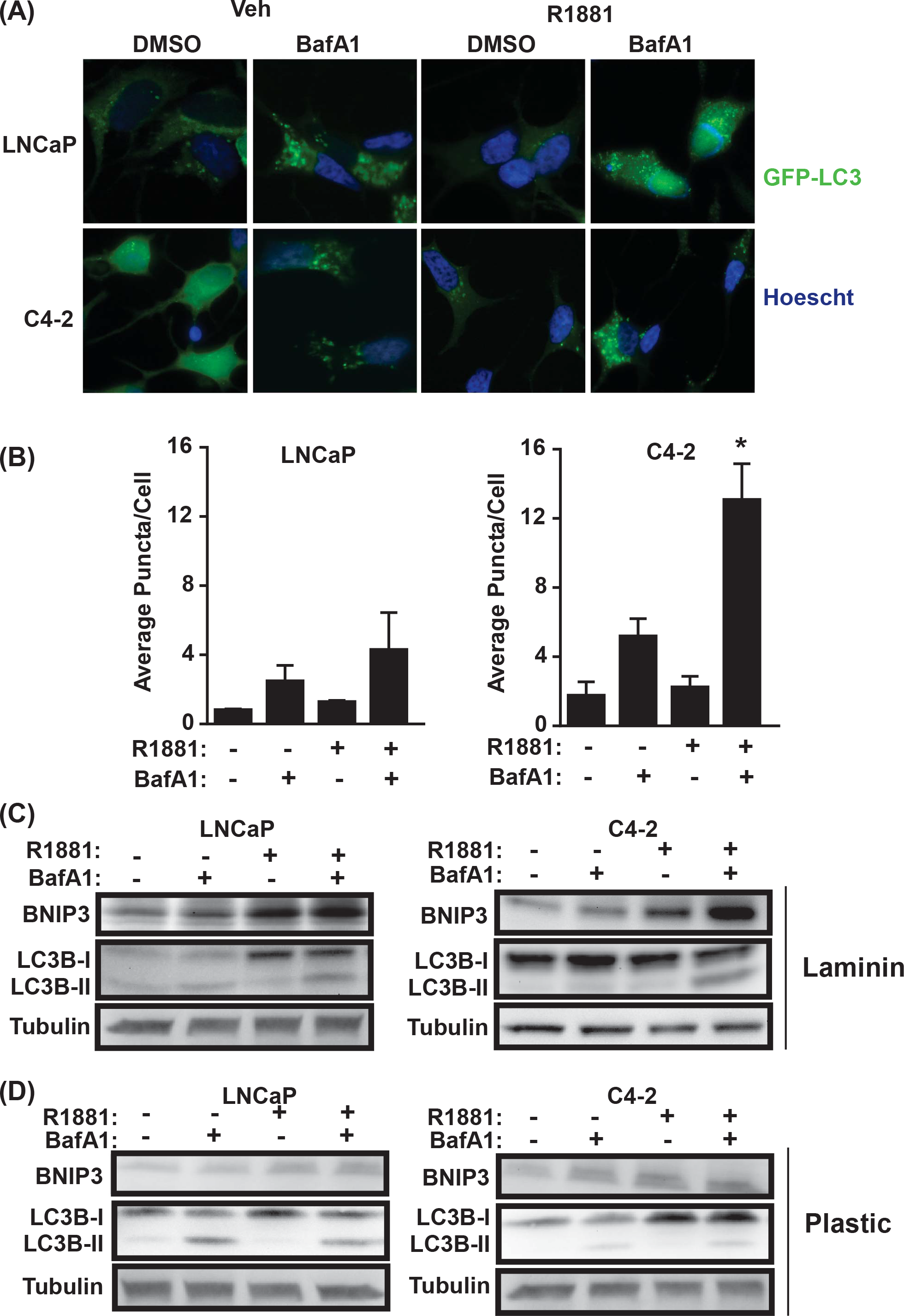
Androgen-induced autophagy requires laminin. **(A)** LNCaP and C4-2 cells stably expressing LC3-GFP and adherent to laminin were stimulated with (+) or without (-) 10 nM R1881 for 24 h and in the absence (-) or presence (+) of 100 ng/mL Bafilomycin A1 (BafA1) during the last 2 h prior to fixation. Cells were stained with Hoescht and imaged by epifluorescence microscopy. **(B)** Quantification of puncta from (A). GFP puncta were counted as positive if the signal was 10 standard deviations greater than background fluorescence. Average is the number of puncta in at least 50 randomly selected individual cells under each condition. * *p*< 0.05, n=3, error bars = SD. **(C)** LNCaP and C4-2 cells adherent to laminin were stimulated with (+) or without (-) 10 nM R1881 for 24 h and in the absence (-) or presence (+) of 100 ng/mL Bafilomycin A1 (BafA1) during the last 2 h. Levels of BNIP3, LC3B-I/II and tubulin were assessed by immunoblotting. **(D)** As in (C), except cells were adherent to plastic. Quantification of immunoblots for (C) and (D) is in Supplementary Figures S2A, 2B.

Because BNIP3 induction requires integrin α6 (see Fig. 2D), we compared androgen-induced autophagy on laminin versus plastic by measuring the conversion of LC3B-I to LC3B-II by immunoblotting. As observed for GFP-LC3B puncta, androgen alone did not strongly induce LC3B-II, but androgen plus bafilomycin resulted in a significant increase in LC3B-II levels specifically in C4-2 cells on laminin (Fig. 5C, Supplementary Fig. S2A). However, combined androgen and bafilomycin did not cause any significant change in the rate of autophagy in either cell line plated on plastic (Fig. 5D). Because BNIP3 binds to LC3B (23,32), it is also degraded upon fusion of the autophagosome with the lysosome. Thus, BNIP3, like LC3B-II, accumulated significantly in C4-2 cells upon combined androgen and bafilomycin treatment (Fig. 5C, Supplementary Fig. S2B). This happened only on laminin and not on plastic (Fig. 5D).

To determine if the androgen-induced autophagic flux occurring on laminin is mediated by integrin α6, cells were treated with integrin α6 shRNA. Blocking integrin α6 expression prevented the accumulation of LC3B-II and BNIP3 in C4-2 cells in the presence of androgen and bafilomycin (Fig. 6A, Supplementary Fig. S2C). To determine if BNIP3 is required for androgen-induced autophagy on laminin, we blocked BNIP3 expression with siRNA and measured LC3B-II accumulation in the presence of androgen and bafilomycin. BNIP3 loss did not significantly affect androgen-induced accumulation of LC3B-II in the presence of bafilomycin (Fig. 6B, Supplementary Fig. S2D).

**Figure 6:**
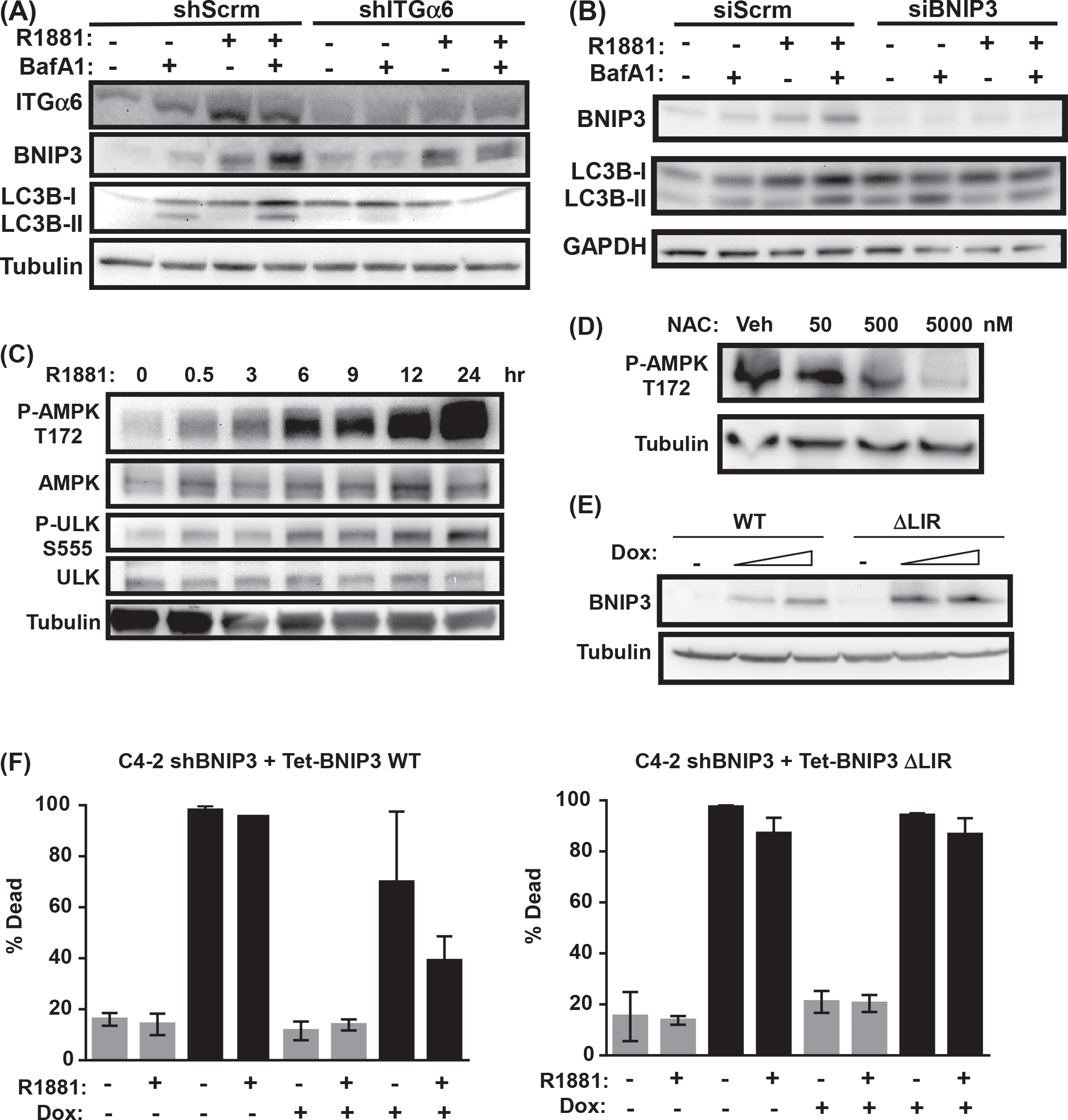
Androgen-induced autophagy requires integrin α6, but not BNIP3, and BNIP3 promotes survival through mitophagy. **(A)** Laminin-adherent C4-2 cells stably expressing an shRNA targeting integrin α6 (shITGα6) or a control scrambled shRNA (shScrm) were stimulated with (+) or without (-) 10 nM R1881 for 24 h and in the absence (-) and presence (+) of 100 ng/mL Bafilomycin A1 (BafA1) during the last 2 h. Levels of BNIP3, LC3B-I/II and tubulin were assessed by immunoblotting. Quantification of immunoblots is in Supplementary Figure S2C. **(B)** Laminin-adherent C4-2 cells transfected with siRNA targeting BNIP3 (siBNIP3) or a control scrambled siRNA (siScrm) were stimulated with (+) or without (-) 10 nM R1881 for 24 h and in the presence of 100 ng/mL Bafilomycin A1 (BafA1) during the last 2 h. Levels of BNIP3, LC3B-I/II and GAPDH were assessed by immunoblotting. Quantification of immunoblots is in Supplementary Figure S2D. **(C)** C4-2 cells adherent to laminin were treated with 10 nM R1881 over a 24 h time course. Levels of activated AMPK (P-AMPK-T172), total AMPK, activated ULK1 (P-ULK-S555), total ULK1 and tubulin were assessed by immunoblotting. **(D)** C4-2 cells adherent to laminin and stimulated with 10 nM R1881 were treated with increasing concentrations of NAC. Levels of activated AMPK (P-AMPK-T172) and tubulin were assessed by immunoblotting. **(E)** C4-2 cells stably expressing shRNA targeting 3’-UTR of BNIP3 (C4-2-shBNIP3) and Tet-inducible WT (Tet-BNIP3 WT) or ΔLIR mutant BNIP3 (Tet-BNIP3 ΔLIR) and adherent to laminin were stimulated with 100 and 1000 ng/mL doxycycline (DOX) for 48 hours and levels of exogenous BNIP3 and tubulin measured by immunoblotting. **(F)** C4-2-shBNIP3 cells expressing Tet-inducible WT (Tet-BNIP3 WT) or ΔLIR mutant BNIP3 (Tet-BNIP3 ΔLIR) and adherent to laminin were stimulated with (+) or without (+) 10 nM R1881 in the presence (+) or absence (-) of 100 ng/mL doxycycline (DOX) for 24 h prior to and in the absence (DMSO) and presence (PX866) of 500 nM PX-866. Cell viability was assessed by trypan blue exclusion 48 hours later. n=3; error bars = SD.

The link between AR and autophagy was also observed in PC3 cells in which constitutively active AR is re-expressed (6). LC3B-II levels were very low upon adhesion of parental PC3 cells to laminin, even in the presence of bafilomycin (Supplementary Fig. S3A). Re-expression of constitutive active AR in two different clones restored LC3B-II levels, which accumulated further upon bafilomycin treatment (Supplementary Fig. S3A). Thus, overall our data indicate that integrin α6-mediated adhesion to laminin and androgen signaling to AR is required to increase autophagy and cause LC3B-II and BNIP3 turnover in the lysosome. Furthermore, these data indicate that while integrin α6 regulates both BNIP3 and autophagy downstream of AR, androgen induces autophagy independently of BNIP3.

### Androgen increases AMPK signaling through ROS

AMPK phosphorylation at Thr172 and activation is induced in response to low levels of ATP in the cell, which is also a trigger for autophagy. AMPK in turn phosphorylates ULK to initiate autophagy (40). Stimulation of C4-2 cells with androgen induces AMPK and ULK phosphorylation within 30 minutes and it continues to increase over 24 hours (Fig. 6C). Increased AMPK and ULK signaling in response to androgen was observed in both castration-resistant lines, but not in parental PC3 cells which lack AR (Supplementary Fig. S3B). Androgen signaling in castration-resistant cells leads to elevated ROS (41), which can stimulate AMPK activation (42). Indeed, treatment of C4-2 cells with the ROS scavenger, N-acetyl cysteine (NAC), decreased androgen-induced AMPK phosphorylation in C4-2 cells (Fig. 6D). Thus, androgen likely acts through the AMPK/ULK sensing pathway to initiate autophagy in response to elevated levels of ROS.

### BNIP3-mediated mitophagy protects CRPC cells from PI3K inhibition

BNIP3 is required for the survival of CRPC cells adherent to laminin and confers resistance to PI3K inhibition, but this is not mediated by BNIP3-induced autophagy. BNIP3 localizes to the mitochondria outer membrane via its C-terminal transmembrane domain with a majority of the protein facing the cytoplasm. Near the N-terminus is an LC3-interacting region (LIR) responsible for bringing mitochondria into the autophagosome for degradation. Mutating the LIR abrogates mitophagy (23,32). To test whether BNIP3 promotes survival through mitophagy, we first generated a C4-2 cell line stably expressing a BNIP3 shRNA targeting the 3’ UTR. We then used a Tet-inducible vector to re-express WT or an LIR mutant (ΔLIR) BNIP3 in the BNIP3-knocked down cells (Fig. 6E). Because endogenous BNIP3 is constitutively suppressed in these cells, androgen could not rescue them from death induced by PX-866 treatment as expected (Fig. 6F). Inducing WT BNIP3 expression with doxycycline protected the cells from PX-866-induced death when androgen was present. However, the ΔLIR mutant prevented the androgen-induced rescue (Fig. 6F). Thus, the pro-mitophagy function of BNIP3 is required to promote survival and protect cells from PI3K inhibition in castration-resistant tumor cells.

## DISCUSSION

There is a marked selection for increased integrin α6β1 expression and decreased integrin β4 expression in primary prostate cancers (5,7,43). Elevated integrin α6β1 is associated with increased invasiveness, lymph node metastasis, and bone metastasis (43,44). We previously demonstrated that AR is responsible for directly inducing integrin α6 transcription and expression in tumor cells (6). In this study, we demonstrate that integrin α6β1 expression is further elevated in castration-resistant prostate cancer. Previous studies, including ours, demonstrated that adhesion to extracellular matrix via integrins confers resistance to many therapeutic regimens, including chemotherapy, radiation, and targeted therapies in many cancer types (45). We demonstrated specifically, that adhesion to laminin via integrin α6β1, confers resistance to PI3K inhibition in Pten-null prostate cancer cells by inducing NF-κB signaling and up-regulating the anti-apoptotic protein Bcl-XL (6). In this study, we identified another integrin α6β1-mediated survival pathway that is selectively operating in castration-resistant prostate cancer (Fig. 7). Enhanced AR activity in castration-resistant tumors leads to enhanced integrin α6β1 expression, such that adhesion to laminin in the presence of androgen activates two distinct signaling pathways. One involves the nuclear translocation of HIF1α, induction of BNIP3 transcription, and localization of BNIP3 on mitochondria. The other involves ROS-mediated activation of the AMPK/ULK energy sensor to enhance autophagosome formation, into which BNIP3-enriched mitochondria are targeted for lysosomal degradation.

**Figure 7:**
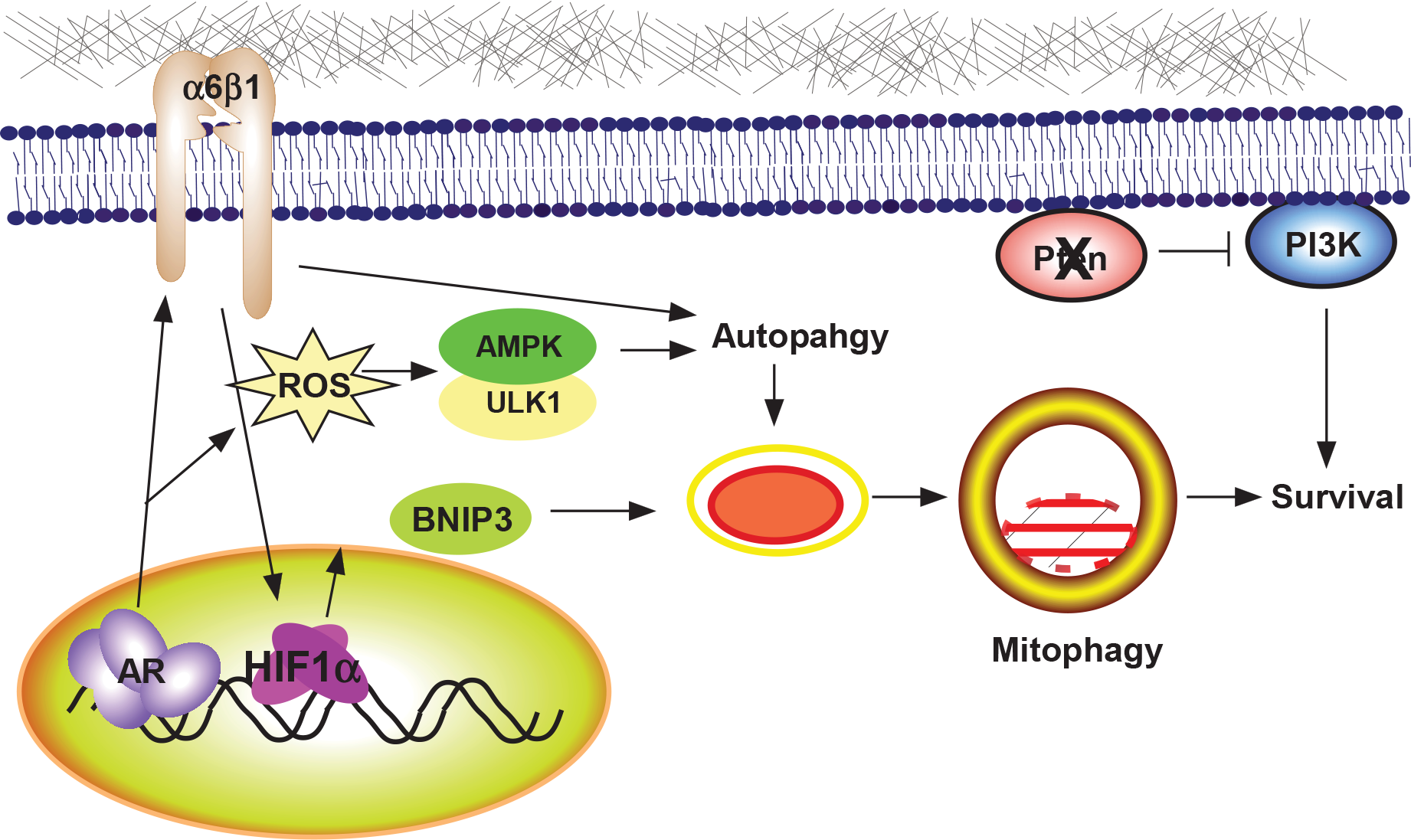
Model for AR-dependent activation of autophagy and mitophagy in castration-resistant prostate cancer through integrin α6β1. Castration-resistant cells harboring active AR and elevated HIF1α stimulate integrin α6β1 expression. Engagement of laminin by integrin α6β1 promotes nuclear translocation of HIF1α and induction of BNIP3. Simultaneously, AR activity induces ROS, triggering elevated AMPK/ULK1 activity, and together with intergrin α6β1 induce assembly of the autophagy machinery. Mitochondria are recruited to the autophagosomes by BNIP3 for targeted degradation. This reduces the level of damaged mitochondria to prevent apoptosis and promote survival. In parallel, constitutive activation of PI3K by Pten loss also promotes survival.

The association of castration-resistance with elevated BNIP3 expression is intriguing and our finding that AR and integrin α6β1 control BNIP3 transcription suggests an interesting mechanism for both acquiring and maintaining a castration-resistant state. Androgen deprivation therapy, particularly in the bone, leads to increased hypoxia (46) and thus elevated HIF1α expression. Recent studies indicate that not only is BNIP3 a direct transcriptional target of HIF1α, so is integrin α6β1 (12,14). Thus, under conditions of hypoxia and no androgen, both integrin α6β1 and BNIP3 are elevated, which promotes survival of tumor cells by limiting ROS production through selective removal of mitochondria. Prostate tumors that amplify AR or gain androgen-independent AR function (through mutation or splice variants), can further sustain and amplify this BNIP3-dependent survival pathway. These castration-resistant tumors may even become adapted to tolerating elevated ROS. In fact, AR increase ROS and ROS induces HIF1α and AMPK signaling (41,47). Thus, ROS production caused by AR signaling, particularly in cells with elevated AR expression, may be responsible for causing the increasing both HIF1a signaling to BNIP3 while simultaneously stimulating autophagy.

Our studies further emphasize the importance of the tumor microenvironment when attempting to understand mechanisms of castration-resistance. Laminin is a major ECM component in lymph nodes and bone, the major sites for prostate cancer metastasis (8,9), and integrin α6β1 is the major integrin expressed on prostate tumors in these tissues (43,44). In cell culture models of prostate cancer, where cells adhere to RGD-containing ECM like fibronection or vitronectin found in serum that coats the plastic petri dish, the effects of laminin-specific integrin signaling are missed. Indeed, we found this AR/integrin α6-induced autophagy and mitophagy pathway to be uniquely activated by laminin in castration-resistant cells. This difference may also explain some of the discrepancies in the literature between our findings and those of others. For instance, a previous report indicated that androgen induces HIF1α signaling through an autocrine loop that depends on PI3K/Akt signaling (48). This suggests that PI3K inhibition would block HIF1α and BNIP3; however, inhibiting PI3K in cells plated on laminin does not prevent androgen from stimulating HIF1α nuclear translocation (not shown) or block BNIP3 expression. Several reports suggested that androgen inhibits autophagy (49,50). A couple of factors may explain this discrepancy. First, those studies failed to measure autophagic flux by blocking lysosomal function, so any effects on increased autophagosome formation are missed if there was also an increase in turnover rate. The importance of this was nicely demonstrated in a recent report where androgen stimulation not only increases autophagic turnover, but also induces the expression of several autophagy genes (51,52). Furthermore, elevated expression of these autophagy genes correlated with poor outcome in clinical cohorts. We also saw a similar increase in autophagy genes upon androgen stimulation (not shown). Second, we were only able to see AR-induced autophagy in the castration-resistant cells and only on laminin, where ROS was required to activate this signal. Thus, the presence of elevated ROS in the castration-resistant cells may help enhance the AR/α6β1-dependent autophagy induction. It also suggests that *in vivo*, where laminin and integrin α6 are abundant, autophagy may remain intact under castration-resistant conditions.

Interestingly, in normal basal prostate epithelial cells that do not express AR, adhesion to laminin via integrin α3β1, is also a strong inducer of autophagy under growth factor starvation (53,54). Furthermore, when basal cells are induced to differentiate into luminal cells, they down regulate integrin expression, up-regulate AR and become dependent on PI3K for survival (6). We also found that death of luminal cells is accompanied by a large increase in BNIP3 (21). We find it striking that the same pathways that are involved in controlling normal prostate epithelial cell physiology become activated and dysregulated in CRPC, as if the cells have a defined set of options with which to evolve and evade selective pressure.

It is well-established that autophagy promotes therapeutic resistance in cancer, and targeting autophagy has provided some promising results in other cancers (55), but has not necessarily translated well to patients. This is complicated by constitutive PI3K signaling in prostate cancer due to loss of Pten or constitutively activated PI3K mutants (4), which suppresses autophagy. Given the high level of PI3K activation in prostate cancer it is surprising that PI3K/mTor inhibitors have not been effective as single agents, or even in combination with anti-androgen therapies (5). Our data indicate that single agent PI3K inhibitors won’t work because AR/α6β1 integrin signaling will still activate Bcl-XL and BNIP3-mediated mitophagy. Furthermore, PI3K inhibition will further stimulate autophagy to augment AR/α6β1 signaling. In combination with anti-androgen therapies, this pathway will still be active due to acquisition of androgen-independent AR signaling and/or gain in hypoxia/ROS signaling. Thus, a multi-prong approach that includes targeting hypoxia and other AR-downstream targets, like integrin α6β1, will likely be necessary to effectively suppress castration resistant disease.

## ACKNOWLEDGEMENTS

We would like to thank Drs. Sander Frank, Eva Corey, and Don Tindall for feedback and constructive suggestions and Penny Berger for technical expertise. We acknowledge the following for important reagents: pBABEpuro GFP-LC3 was a gift from Jayanta Debnath (30), pLenti CMV rtTA3 Blast (w756-1) (Addgene plasmid # 22405) and pLenti CMV Neo DEST (705-1) (Addgene plasmid # 26429) were gifts from Eric Campeau (33).

These studies were supported by funding from NIH/NCI CA154835 (CKM, EAN, VSS) and the Van Andel Research Institute. Additional support was provided by NIH/NCI CA159406 (AEC).

